# The measurement of species selection on evolving characters

**DOI:** 10.1101/176438

**Authors:** Carl Simpson

## Abstract

Many processes can contribute to macroevolutionary change. This fact is the source of the wide variety of macroevolutionary change across time and taxa as well as the bane of pale-obiological research trying to understand how macroevolution works. Here, I present a general framework for understanding the variety of macroevolutionary phenomena. Based on Price’s theorem, this framework provides a simple quantitative means to understand (1) the macroevolutionary processes that are possible and (2) the way those processes interact with each other. The major qualitative features of macroevolution depend first on the number of processes that co-occur and then on the magnitudes and evolutionary directions of those processes. Species selection, the major macroevolutionary process, consists of patterns of differential rates of speciation and extinction. Its macroevolutionary efficacy depends on the presences of sufficient microevolutionary change. Conversely, microevolutionary change is limited in power by the independent evolution of species, and species selection acting across populations of species can amplify or suppress microevolution. Non-trends may result if species selection sufficiently neutralizes microevolution and may yield stable macroevolutionary patterns over many millions of years.

## Emergence is a fact of life

Due to the cumulative addition of new levels of organization, life has an inherently hierarchical structure (Maynard Smith and Szathmry 1995, McShea 2001a, b, McShea and Simpson 2011, Michod 1999, Simpson 2011). This structure is obvious in the bodies of multicellular and colonial animals, as they consist of cells and animals that were independent entities in earlier evolutionary times. What is less obvious is that life’s hierarchical structure does not stop at the boundaries of bodies (Simpson 2011, Van Valen 1980, Van Valen 1991). It continues to even more inclusive levels of populations (colonies after all are homologous to, and derived from, populations) and species. Because populations are structured spatially, temporally, and genetically, the details of how they are structured must be relevant to understanding the evolution of, and the levels of selection acting at, each of these more diffuse levels of organization.

Among types and levels of populations, the species level stands out as special (Gould 2002, Stanley 1975), independently of your preferred species concept. This is because it represents the most inclusive and cohesively (stable) structure that populations have attained. This stability may be produced in many different ways (related to species concepts) from a shared genetic pool, reproductive compatibility, stable morphological similarity, shared evolutionary history, common ecology, etc. Even if these factors yield species with fuzzy bounds, they add coherence to the species level. The ultimate evidence for the coherence of species is the fact that speciation can lead to new independent species and that extinction can lead to their permanent end.

As argued in the companion paper (Simpson 2016), speciation and extinction gift species with the ability to generate a new level of selection that is independent (screened off, in the sense of Brandon 1982, Sober 1992) of, and emergent from, lower levels of selection, such as the organism, colony or group, or cellular levels of selection. The components of fitness at the species level are determined by speciation and extinction, whereas the components of fitness at the organismal level are determined in part by the birth and death of organisms. In the companion paper (Simpson 2016) I argue for and provide evidence of the independence of speciation and extinction from organismal birth and death. For example, there is no evidence for a correlation between an increase in organismal-level fitness (as expected over the duration of a species) and an increase in speciation rates over the same timescales. This observation rejects the effect-hypothesis (Vrba 1983, 1984) as an explanation for species-level selection. Similarly, extinction happens to all species, often after significant timespans where microevolution is rampant. This means that even increasing fitness from the microevolution of organisms within species does nothing to prevent extinction. This is the specific insight that Van Valen had when he proposed the Red Queen’s hypothesis (Van Valen 1973)—no amount of microevolution and therefore fitness increase can prevent extinction. The proposed mechanism for Van Valen’s Red Queen’s hypothesis comes from higher-level processes where the interaction among species, rather than the interaction of organisms with their environment, leads species to extinction.

At the macroevolutionary scale, patterns of speciation, extinction, anagenetic change, and cladogenetic change can all interact to produce large-scale trends. Although we have known of the importance of all four of these factors to macroevolution for sometime (Alroy 2000, Eldredge and Gould 1972, Gould 2002, Gould and Eldredge 1988, Gould and Lloyd 1998, Grantham 1995, Jablonski 2008, Lieberman et al. 1993, McShea 1994, Stanley 1975, Stanley 1979, Turner 2010, Wagner 1996), there has been little progress in understanding how these process can interact. This understanding is important because all four processes can occur together in nature–Not only may species selection occur, but stasis is common but it is not ubiquitous (Hopkins and Lidgard 2012, Hunt 2007, Hunt et al. 2015) and therefore anagenesis occurs, and cladogenetic changes between species may be common (Hopkins 2016). Any general framework of macroevolution must be able to understand all possible cases. In this paper, I provide a general framework for understanding the interactions among macroevolutionary processes in order to understand how different interactions can yield specific observable macroevolutionary patterns. I will propose a means to distinguish between species selection and drift, how anagenetic and cladogenetic change interact with species selection to modulate and steer largescale trends, and show how this framework can be generalized to understand macroevolutionary change among multiple covarying characters.

## Operationalizing macroevolution

Now that we know to look directly for patterns of differential diversification in macroevolution rather than use indirect inferences (Simpson 2016), we can move forward and study how differential diversification came to be and what its macroevolutionary consequences may be. The cause of differential diversification maybe complicated, and involve hitchhiking between characters (Levinton 2001, Wagner 1996) and it can also be simply random drift with no real cause at all (Chevin 2016, Gould 2002). These topics have been largely unexplored in macroevolution because understanding the basic features of species selection have been much more pressing.

The way forward was pointed out in Arnold and Fistrup’s (1982) paper on the hierarchical expansion of evolutionary theory. They introduced Price’s theorem to paleobiology, which they used to delineate hierarchical levels and also to define the emergent fitness approach. Their partitioning of levels matches the way I have outlined elsewhere (Simpson 2010, 2011, 2013, Simpson and Müller 2012). The arguments I presented in the companion paper (Simpson 2016) serve to justify the emergent fitness (Arnold and Fristrup 1982, Gould 2002, Grantham 1995) approach over the alternative emergent trait approach (Lieberman et al. 1993, Lieberman and Vrba 1995, Vrba 1983, 1984) as having better empirical fit to what we can observe in nature.

More recently, I have used Price’s theorem to understand how micro- and macroevolutionary processes interact (Simpson 2010, 2013, Simpson and Müller 2012). This approach, is derived from a general form of quantitative genetics (a field that focuses on phenotypic evolution), and is ideal for macroevolutionary study because it permits us to understand how many processes interact to produce a macroevolutionary result. The terms of Price’s theorem are also simple statistical signals that we can easily measure empirically. Price’s theorem serves both as a guide for analysis and a tool for synthesizing all macroevolutionary processes.

Price’s theorem works because it identifies why the mean trait value of a population changes a particular way over time. It does this by partitioning the change in trait values into components—the different ways that the mean of a distribution can change—and summarizes how these ways interact with each other to make change happen.

There are three ways to change mean trait value of a population in terms of how the members reproduce and change over time. First, members with particular trait values may diversify at different rates. The pattern of covariation between diversification potential and trait values determines what the relative frequencies of members with particular traits over time. This covariance between diversification rate and trait values is the essence of selection. And selection shares this same statistical signal no matter the level at which it is operating.

Second, descendants may differ from their ancestors in some way. The similarities between these two generations can be summarized by their covariance as well. This pattern of similarity between ancestors and descendants encapsulates the cladogenetic changes that punctuated equilibrium first recognized. For example, if Wright’s rule, a pattern of unbiased change between ancestors and descendants (Eldredge and Gould 1972), occurs then the covariation between ancestors and descendants will be near one. Melanie Hopkins recently showed that there is a wide range of variation in the magnitude and direction of speciation that may or may not be constant with Wright’s rule (Hopkins 2016).

Finally, the traits of members within a population can change over time. This anagenetic change is summarized by the average change of members over time. For example, this term summarizes any tendency for a member to increase in size over time. Morphological stasis is a special case, where the average change is nearly equal to zero. An average change of near zero is also possible if the way traits change is similar to an unbiased random walk or if there are diverging patterns of directional change (as in the hypothetical rabbit example in Simpson 2016).

Price’s theorem simplifies the complex conceptual issues and empirical observations that make up macroevolution. I believe it serves as the foundation of a formal macroevolution theory and as a tool to ask new questions.

## Measuring species selection

It is easy to measure species selection empirically as the covariance between net diversification rate and trait values. Alternatively, diversification rate can be partitioned into speciation or extinction rates and their covariance with trait values defines how much of species selection is driven by differential speciation or extinction.

The covariance simplifies things analytically and conceptually because the covariance can be broken down into two constituents, the linear regression between emergent fitness and traits and the variance of the traits. This means that we can capture directional species selection simply with a linear regression of diversification rates and trait values (Fig. 1).

**Figure 1:**
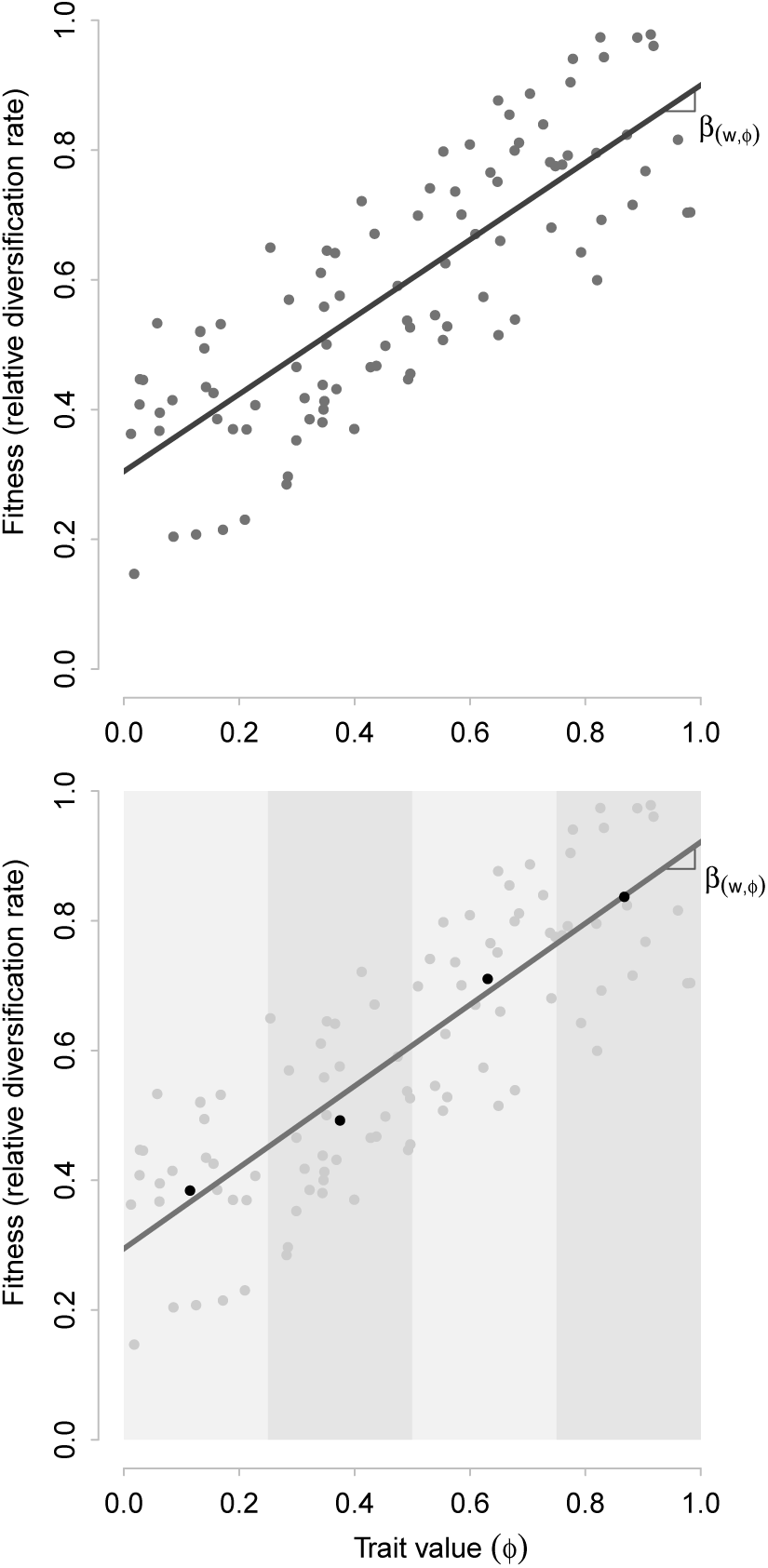
The linear regression of diversification rate on trait value measures species selection. Ideally (top panel) speciation and extinction rate is measured for each species. The slope of the linear regression though these points is the selection vector and predicts how much and in what direction the population will shift morphologically over time due to differential diversification. In typical fossil data (bottom panel), speciation and extinction rates can only be estimated from the distribution of rates observed within a set of species. A linear regression through single rate estimates for species binned by trait values (here denoted by alternating grey bars) yields a measure of species selection comparable to the ideal case.

The linear regression of diversification rate and trait values is the magnitude and direction of species selection. A positive regression indicates that species selection will increase the value of a trait over time. A negative value means that trait values will decline. The slope of the regression is equal to how much the trait values are expected to change by.

Importantly, there is a special case, where the slope is equal to zero. This is the case where there is no directional species selection. In this case, there is no differential diversification to induce a change in the average trait values over time. If this occurs, then the only process that can drive trends in the mean are microevolutionary.

However, more complex selective scenarios, for example stabilizing or disruptive species selection, can also be analyzed with this framework. Just as their organism-level analogs do, these macroevolutionary processes change the amount of variation in a trait is expressed within the population of species. Stabilizing species selection decreases the amount of variation and disrupting species selection increases it. To make a calculation of the amount of stabilizing or disruptive species selection, the trait values simply need to be transformed to measure their dispersion from the mean value by taking the squared deviation of a species’ trait from the mean of the population (Lande and Arnold 1983, Rice 2004). The covariance between fitness and the deviation measures stabilizing and disruptive selection. A positive covariance equals disruptive selection because members at the extremes (greater deviation from the mean) have a higher diversification rate than members near the mean. A negative covariance indicates stabilizing species selection because the members near the mean have the highest relative diversification rate.

The one empirical challenge in measuring species selection stems from how we measure diversification rates. Ideally we would measure the expected diversification rate of each species individually. This estimate could vary continuously and its covariance with continuous trait values would be easy to estimate. A species-specific diversification rate may be possible to estimate with a phylogeny or with machine learning (Finnegan et al. 2015), but otherwise we must lump species together in some way to estimate species selection. We can lump species together by trait values to estimate discrete diversification rates for each pool of species defined by possessing similar trait values (Simpson 2010, 2013, Simpson and Harnik 2009, Simpson and Müller 2012). With this type of lumping we then perform a linear regression to estimate species selection (Fig. 1). Alternatively, we can lump species together by their survival or recent speciation and perform a logistic regression that preserves continuous trait variation, but discretizes diversification (Finnegan et al. 2008, Harnik et al. 2012, Payne and Finnegan 2007).

## Microevolutionary change

With the appropriate samples, the magnitude of anagenetic change can be directly observed in the fossil record or from historical time series (Cheetham 1986, Gingerich 1983, 1993, Hendry and Kinnison 2001, Hendry and Kinnison 1999, Hopkins and Lidgard 2012, Hunt 2007, Hunt et al. 2015, Kinnison and Hendry 2001, Uyeda et al. 2011). Cladogenesis can also be directly measured if there is a phylogenetic framework that defines the relationships between species (Bokma 2008, Hopkins 2016, Hunt 2013, Ingram 2011, Simpson 2013). However, when analyzing anagenesis and cladogenesis in a macroevolutionary scenario, their patterns need to be summarized by the change in phenotype from time interval to time interval rather than as a statistical summary of their mode. In terms of Price’s theorem, both anagenesis and cladogenesis contribute to two parameters in Price’s theorem. First, the phenotypic variation is the result of the variation of traits across the population of species. The variation in a single interval of time is the result of prior patterns of species selection, anagenesis, and cladogenesis.

The second parameter measures the change due to anagenesis and cladogenesis additively interacts with the selection covariance and is solely defined by the patterns of change due to anagenesis and cladogenesis from interval to interval. In the framework of Price’s theorem, it does not matter if anagenetic change is directional, random, or shows stasis because any change, no matter how small, will play a part in influencing a change in the mean trait value in the population of the species. The average change, calculated across all species, is what is relevant.

## Micro and macro interact

In traditional macroevolutionary thinking, the average anagenetic and cladogenetic change interact only additively with species selection (Gould and Eldredge 1977). This additive interaction, does occur as described above, but there is also an additional interactive affect. Price’s theorem has another parameter, the heritability between ancestral and descendant species, that is caused by anagenesis and cladogenesis directly. Like the standing variation in traits, the heritability interacts multiplicatively with species selection. Through heritability, the pattern of microevolution modulates the response to species selection.

Heritability can be defined narrowly or broadly. The narrow definition is the amount of additive genetic variance. Narrow sense heritability is useful for extremely small scale microevolutionary situations, such as breeding. In this scenario, the additive genetic variance gives us a measure of the similarity between ancestors and descendants that excludes sources of variation that breeders want to factor out, such as environmental influences, or complex developmental interactions. Heritability in the broad sense measures only the covariance in form between ancestor and descendants. And so it incorporates all mechanisms for similarity. For our purposes (at the species level) broad sense heritability is the only useful option because we are not yet interested in (or there may not be) a genetic component to how traits, such as geographic range size, come to be. At the species level, heritability measures the similarity between species in adjacent time intervals. The population of species in the subsequent time interval consists of some species that survived and some new species that are descendants of ancestors from the previous interval. These two subpopulations also delimit the anagenetic and cladogenetic components of heritability. The anagenetic component measures the similarity between successive morphologies of the same species that persist over time. A species in morphological stasis will have the same morphology (or nearly the same) in both time intervals and will have a heritability near 1. But a species evolving directionally will be different in the two intervals and that evolutionary change will consequently lead to a different heritability. The cladogenetic component measures the similarity between the ancestor in one time interval and its descendant species in the subsequent interval.

The estimation of anagenetic heritability requires repeated observations of traits in multiple time intervals. We can then measure the heritability as the linear regression between the traits of anagenetic species in adjacent intervals (Fig. 2). To directly estimate the cladogenetic component of heritability it is important to have a phylogenetic framework that identifies ancestor-descendant relationships. With this information, it is easy to plot the descendent trait value against the ancestral value and calculate the linear regression to estimate this component of heritability.

**Figure 2:**
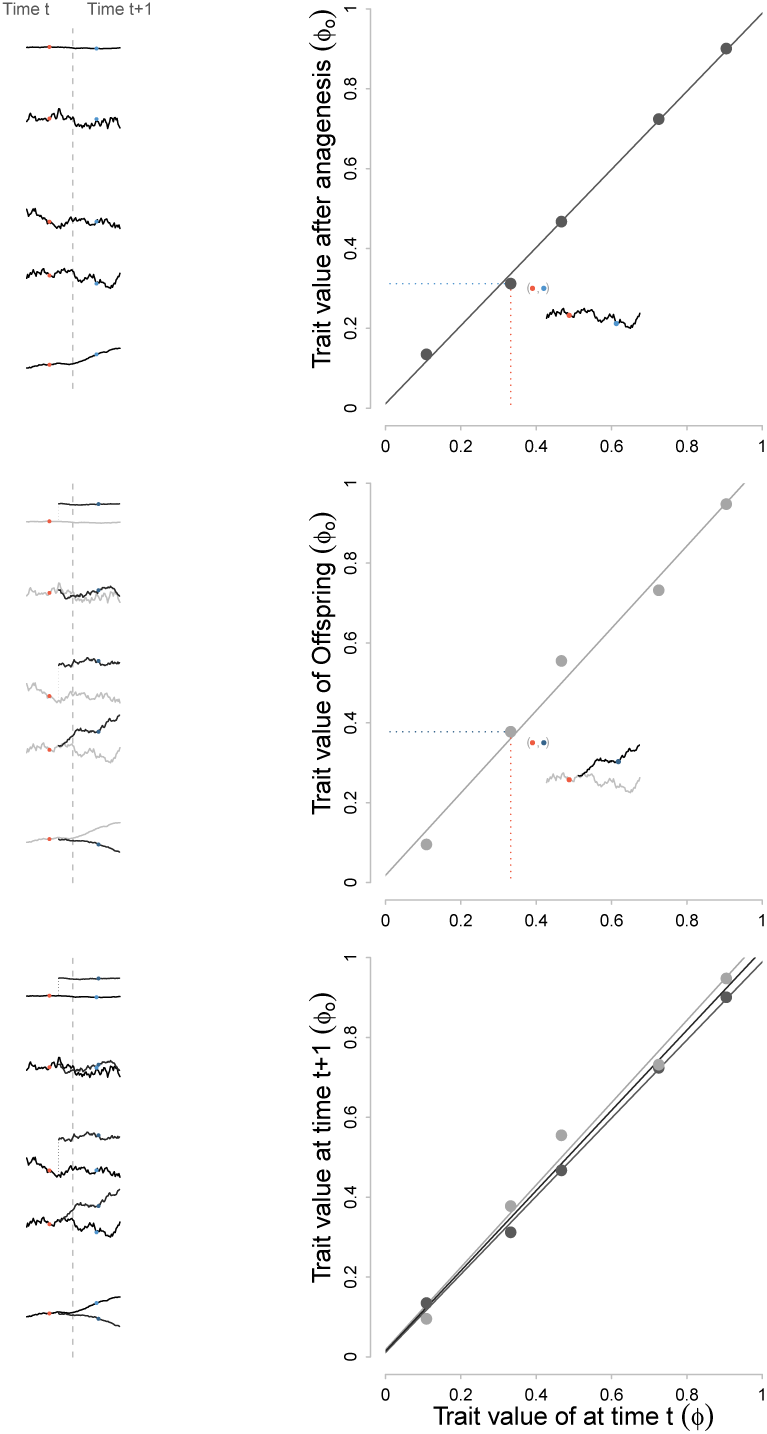
The linear regression of traits for species in adjacent intervals of time measures the heritability of traits. There are two contributors to the pool of species in adjacent intervals of times. Species can survive from one interval to the next and the heritability of their anagenetic change can be measured by regression species at time t+1 onto species at time t (top panel). New species can occur in the second interval that descend from ancestral species present in the first interval. Regressing descendent species onto ancestral species cladogenetic measures the heritability of cladogenetic trait change (middle panel). The total heritability consists of anagenetic and cladogenetic components (bottom panel). The total heritability is required for macroevolutionary analysis and can be calculated as the linear regression through all anagenetic and cladogenetic ancestor-descendent comparisons or as the average of anagenetic and cladogenetic regressions. The evolutionary trajectories in trait values for several species are shown in the left column. The sampled trait values are in adjacent bins are shown in red and blue and are used calculate the heritability.

There are two ways to estimate the total heritability that incorporates both anagenetic and cladogenetic components. First is to take the linear regression of the pooled anagenetic and cladogenetic comparisons. Equivalently, the average of the anagenetic and cladogenetic regressions will give the total heritability.

The sign and slope of the heritability regression is informative about the tempo and mode of microevolution. For the total heritability, a regression of 1 signifies the classic punctuated equilibrium scenario, with morphological stasis and Wright’s rule. In this case, species and stay the same across adjacent intervals and the direction of speciation is random. A slope greater than one means species are on average increasing trait values and a slope less then one means that species are decreasing in their trait values. For certain traits, it may even be possible to have a negative heritability. This scenario could occur if species tend to differ from their ancestors in the trait especially if the trait has limited numbers of states.

## Species selection, hitchhiking, and drift

### Species selection

Many arguments about levels of selection highlight the scenario where selection at two levels oppose each other (Gould 2002, Rice 1995, Slatkin 1981). This scenario was seen as providing the best chance of demonstrating the efficacy of species selection—only if we could demonstrate empirically that species selection could oppose microevolutionary change would we have conclusive evidence that species selection is a real and effective process. Yet very few examples of opposing selective process has been identified. The first examples are Van Valen’s (1975) identification that microevolutionary increases in mammal body size is topped by selective extinction of large sizes and results in an equilibrium body size distribution. In a more recent example, Goldberg and others (2010) found that self-incompatability in the nightshades plant family is maintained by species selection despite the short-term advantages of self-fertilization at the organismal level.

What the multiplicative interaction between species selection, cladogenesis, and anagenesis provides is a dramatically different view on the interaction macroevolutionary and microevolutionary processes. With this approach, we have a tool that allows a detailed dissection of the evolutionary processes involved in any empirical case, regardless of the relative orientation of species selection and microevolution. The multiplicative effects between micro and macroevolution are particularly important in the evolution of multiple covarying traits. But even in the evolution of single traits like body size or self-incompatability, there can be a strong effect.

The worst case scenario for species selection under the old paradigm must be the situation where species selection and microevolution operate in the same direction. With the help of Price’s theorem, this situation is actually a useful case for showing the efficacy of species selection. The change in the average trait value is the product of species selection, trait variance across species, and the heritability of traits from ancestor to descendent species. Both the variance and the heritability across species are produced exclusively by microevolutionary changes. If there is trait variation across species and all evolutionary process are operating in the same direction, species selection will amplify the trend, making it stronger than microevolution alone could produce, proportional to selection’s magnitude.

There is major conceptual significance in the multiplicative interaction between species selection and microevolution (Fig. 3). It means that macroevolution requires microevolution to work. In the absence of microevolution change and variation, species selection cannot produce trends. But in the absence of species selection, microevolution can produce trends alone. With the absence of species selection the magnitude of a microevolutionary trend is no longer a function of the variation among species as it is with species selection. The dependence of the magnitude of trends involving species selection with amount of variation could be a useful and easy tool to indirectly survey for the occurrence of species selection.

**Figure 3:**
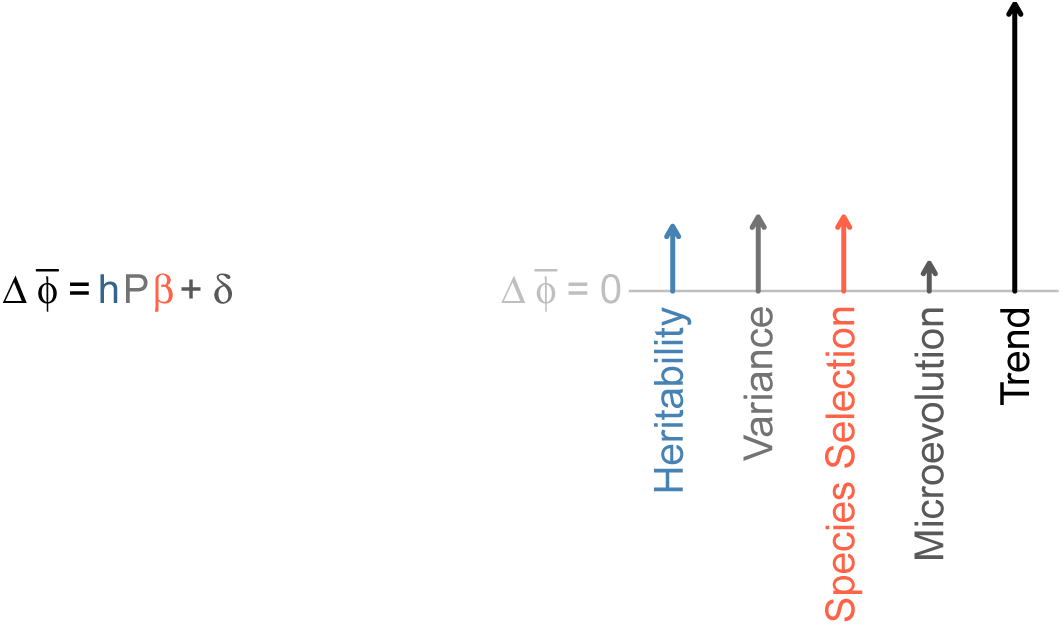
Macroevolutionary processes interact according to Price’s theorem Here a general form of this theorem tracks the change in mean traits values 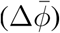 due to the product of heritability (h), phenotypic variance (P), and the selection vector (*β*) plus the average microevolutionary change due to anagenesis and cladogenesis (*δ*). The right panel shows visually how the magnitude and direction of these processes interact. The multiplicative term determines the majority of the magnitude and direction of the trend. The consequences of varying the magnitude and direction of parameter in turn is explored using the same visual vector field in Figure 6.

### Multiple traits and hitchhiking

Selection, whether at the organismal or at the species level, rarely acts on single characters alone because organisms and species possess a number of characteristics. It may be common for characters to covary with each other as well. At the species level, the dimensions of rarity (geographic range size, abundance, and niche breadth) covary with each other (Harnik et al. 2012). Diverse types of traits can also covary and influence extinction or speciation rates (Harnik 2011). Understanding the covariance between characters is important to understanding the response to species selection and also its causal structure. With rarity, once the covariance among its dimensions are accounted for, only geographic range size showed consistent extinction selectivity (Harnik et al. 2012).

With multiple traits, the structure of the trait covariance matrix provides an empirical means of understanding the causal links between traits and fitness. Using the logic of path analysis (Wright 1920, 1960), we can build a trait covariance matrix that incorporates hypotheses of causal structure. With path analysis, the covariance between a trait and fitness is calculated by summing the covariation along the paths from trait to fitness in a causal map. The causal map consists of partial regression coefficients between traits caused by others in the map and covariances between source traits that are not influenced by other traits in the map. We can empirically test different causal maps to find the most likely interactions between traits (Harnik 2011). In the case of a hitchhiking trait, its covariance with fitness will be the sum of all the paths between it and fitness, only via its connection with linking traits.

Interacting traits complicate the trajectories that macroevolution can take, as it depends not only on the selection on each trait and each trait’s variation, but the covariation among all traits as well. The result of all this interaction is a macroevolutionary trajectory that is a compromise between the patterns of selection on each trait. Figure 4 shows how interactions between traits cause deviations from the path of selection. In the univariate cause, the pattern of selection largely determines the direction of the trend. But with two covarying traits the direction of the trend is not always in the same direction as the direction of selection on either trait alone. This deviation is most extreme in the case of hitchhiking, where a trend can occur in a trait that only interacts with selection though a covarying intermediary trait.

**Figure 4:**
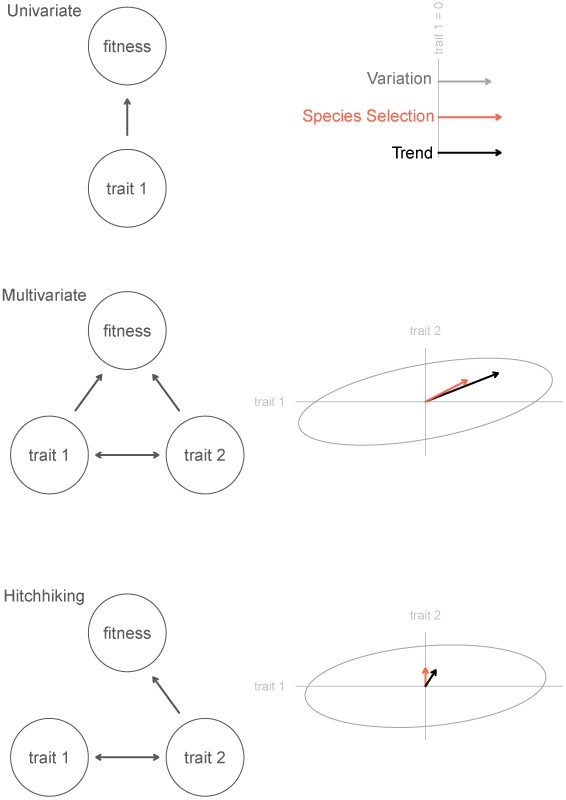
The interactions between traits and fitness have consequences for the magnitude and direction of their evolution. The interactions between correlated traits and differential diversification can be studied using path diagrams. These three examples show selection on a single trait (top), selection of two covarying traits (middle), and the hitchhiking of one trait on another with a direct covariance with fitness (bottom). The resultant evolutionary change for selection on each trait is shown, assuming a perfect heritability and no microevolution change. In the univariate case, the trend follows the direction of selection increasing or decreasing the mean trait value promotional to its variance. In the multivariate case, the direction of a trend may deviate from the direction of selection depending on the covariance between traits and the relative magnitude of the partial regression of fitness on each trait. For hitchhiking, the partial regression of fitness on the hitchhiking trait is equal to zero, but there is still a directional component to its evolutionary change due to the traits covariance with an intermediary trait that directly covaries with fitness. The causal structure implied by path diagrams can be used to test between causal hypotheses and to build a phenotypic covariance matrix (P in Figure 3) for macroevolutionary analysis.

### Drift

When studying species selection we want to know how much of the pattern of differential diversification that we see is a consequence of an underlying causal structure. Because the causes of extinction and speciation can be so complex, traits values may not completely determine fitness. The uncaused or stochastic variation in fitness is termed drift. By chance, even random variation in fitness across species can have a non-zero regression with trait variants and so mimic selection. Drift is not an empirical nuisance, rather it is a real process that has evolutionary consequences. Any non-zero regression caused by drift is enough to add a directional component to a trend and result in an important macroevolutionary change.

Because drift is stochastic, we don’t expect the trend to be maintained over time. And so, over long time frames, drift won’t generate trends, just as on average, it is unlikely to toss a long streak of heads in a large sequence of coin tosses. But just as there can be runs of heads or tails in a series of coin tosses, drift in any actual instance may be a common contributor to trends. Drifts contribution to differential diversification may amplify or dampen trends at any time.

I have been tempted to think of drift in terms of the temporal patterns of its resulting trend. With this line of thought, drift would be a random walk in the mean trait value over time. Drift would cause a trend that changes direction and magnitude randomly. The problem with this view is that selection can also change direction and magnitude (Grant and Grant 2002). By comparing only the temporal resultant of selection or drift, it can be very difficult to distinguish them because we are trying to distinguish between two processes both of which produce the same trend pattern. A better alternative is to find another pattern that uniquely distinguishes between selection and drift at the point where their distinction occurs.

Luckily, Sean Rice’s recent work extending Price’s theorem to include stochastic processes (Rice 2004, 2008, Rice and Papadopoulos 2009, Rice et al. 2011) gives insights on partitioning the effects of selection and drift. Using some of his tools, we can distinguish selection and drift, without adding significantly to the types and amount of data that we need to collect to study species selection. Rice’s insight stems from considering the fitness of any unit as a random variable. In terms of species selection, this means that for any species possessing a particular trait, its emergent fitness is not fixed but instead is a random variable pulled from a distribution of possible fitness values.

Consider as an example extinction selectivity on the geographic range sizes of snail species. Snail species with small geographic range sizes have relatively higher extinction rates than snail species with larger ranges. The relative magnitude of the extinction rates among range size classes could be determined by a chain of causation linking the size of ranges to survival, in which case the differential would be selection. Alternatively it could be stochastic and the differential in extinction would be drift. We can’t distinguish between selection and drift by comparing the differential pattern of selectivity alone because selection and drift can both produce the same differential pattern. Instead, our ability to measure selection and drift requires us to know the amount of variation in extinction rate among small ranging (and separately for large ranging) snail species. If this variation is small, so that all small ranging snail species have similar extinction rates, then there is little opportunity for drift to operate because a species all have similar extinction rates. If however the variation in extinction rates is wide among small ranging snail species, then drift plays a part.

As an illustration of drift, we can imagine subsampling species (within a size class) randomly and comparing their extinction rates. When drift is strong, a random subsample of species will yield inconsistent rates and each addition subsample of species would give a different pattern of extinction rates. When drift is weak, different subsamples of species will not differ much in their extinction rates. The more directly a trait influences speciation or extinction, the less speciation and extinction will vary among species with the same traits. The more important drift is, the more species with the same traits will vary in speciation and extinction rates. The importance of drift is proportional to the variation in fitness of species with similar traits.

Even with drift, and variation among species with the same trait, selection can still occur if the fitness among species of different traits do not overlap significantly. Figure 5 illustrates how drift and selection can occur together by the interplay between variation in fitness within traits and differences in mean fitness among traits.

**Figure 5:**
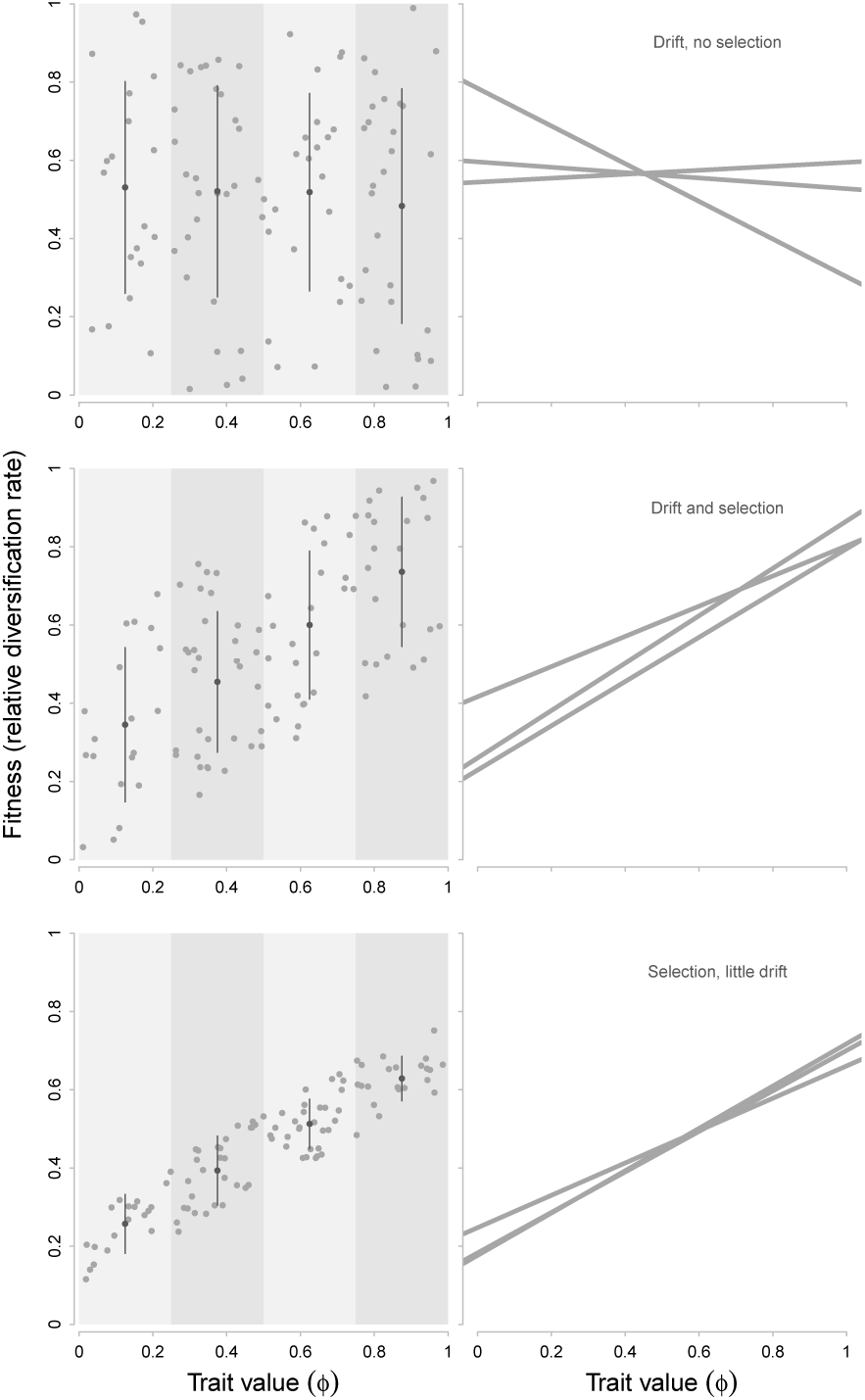
Drift at the species level can mimic species selection by introducing stochastic directional change. The potential for drift to influence trends is always present, but its effectiveness is dependent on the variation in fitness across species with the same trait value. The three panels in the left column show the pattern of net diversification rates for species with similar traits (four traits binned by the grey bars). The dot and whiskers plotted in each grey bar shows variation in fitness for these species, with the width of the bars showing plus or minus one standard deviation. The consequence of drift is shown in the panels on the right column, where the selection differential (linear regression of fitness on trait values) is shown for a random subset of the species pulled from the plot shown in the adjacent left column. The effect of drift is shown how variable the slopes can be. As the variation in fitness of species of each trait declines, the effect of drift also declines. Selection and drift can cooccur because drift is a consequence of the variation in fitness for species with the same trait and selection is a consequence of the variation in fitness for species with different traits.

## Trends and non-trends

As we have seen, there are a large number of macroevolutionary processes. Some arise, like anagenesis and cladogenesis, from microevolution. Species selection, drift, and hitchhiking are emergent macroevolutionary processes. With the help of Price’s theorem, we can understand how these processes interact with each other to result in a change in the average trait value of a clade. The multiplicative terms in Price’s theorem are particularly important because it shows that macroevolution depends on microevolution to operate. If microevolution does not produce any variation among species, then no matter the strength of species selection there can be no macroevolutionary change (Fig. 6).

**Figure 6:**
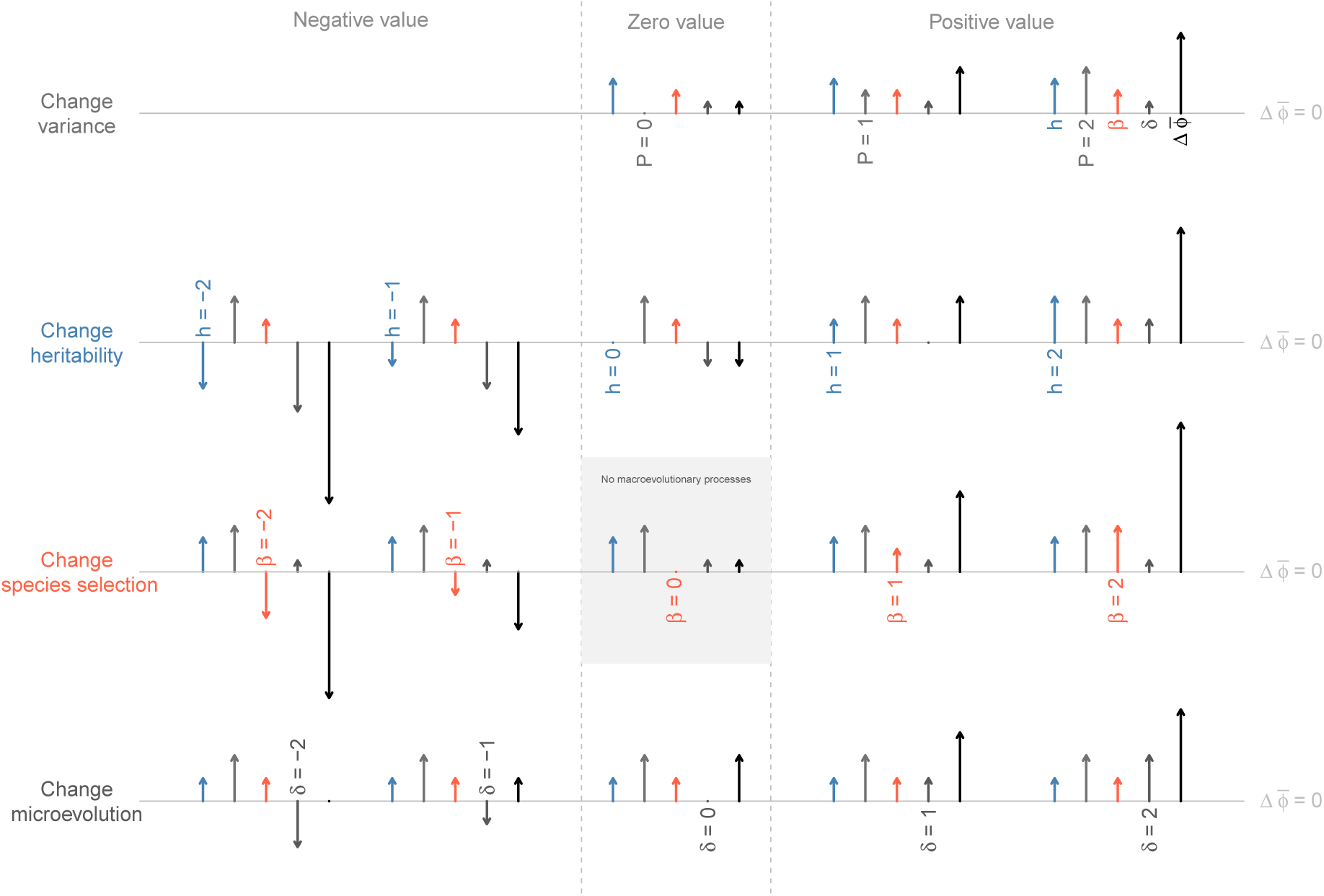
The importance of macroevolutionary and microevolutionary processes on trends is described using visual vector fields defined in Figure 5. Each row shows the effect on a trend of varying a single species-level parameter. The magnitude and direction of that parameter changes horizontally, from strongly negative on the far left, weakly negative on the middle left, a value of zero in the center, weakly positive on the middle right, and strongly positive on the far right. Each horizontal line defines a trend of zero net change. Vectors that point up are positive and those that point down are negative. In the top row, species-level variance can not be negative, so trends are not possible in that left region. The central grey rectangle shows the only parameter combination here without species selection. In this case, the trend is solely generated by the average pattern of anagenesis and cladogenesis and is independent of the amount of species-level phenotypic variation within the clade at any one time.

But there are other interactions between micro and macroevolution that also result in no net trend that are especially common when there is selection on multiple covarying traits. In this case, if the covariation among traits is orthogonal to the direction of species selection the result is no net trend. Selection pulls the population in one direction, but covariation among traits (which is produced by microevolution) constrains the population of species. The interaction between trait variation and selection fixes the mean trait values in a position that would not occur if species selection or microevolution acted alone. I call the patterns resulting from this scenario non-trends.

Non-trends can be more stable than a purely passive trend because of the tension between processes maintains a stable trait distribution. A passive trend (in the sense of Stanley 1973, McShea 1994 and Wagner 1996) occurs by stochastic variation in anagenesis, cladogenesis, and species drift such that variation expands out and the mean trait may shift. Non-trends on the other hand are stable because two or more opposing directional evolutionary processes work against each other, limiting the opportunity for directional change at the same time as limiting the opportunity for variation among species to accumulate.

Non-trends produce the same pattern that the Ornstein-Uhlenbeck (O-U) model of stochastic microevolutionary trait change produces, but by a very different mechanism. The O-U process does not model any hierarchical structure in macroevolutionary processes. Rather, it defines an adaptive peak that lineages evolve towards through anagenesis and cladogenesis. Variation in the direction of these microevolutionary rates leads the clade toward the adaptive peak and keeps it near by once it evolves to occupy that peak. Non-trends reach a similar state of stability from the interaction between species selection and microevolutionary change as those two processes find an equilibrium. And so, the equilibrium point of a non-trend is resembles the adaptive peak of an O-U process in that it produces a stable average trait value in the clade. I suspect that a significant number of results empirically identifying O-U processes will in fact be examples of non-trends if we where to go back and look for evidence of species selection.

There are three documented examples of non-trends in the literature. The two oldest are Van Valen’s (1975) examples in foraminifera and mammals, where species selection is opposed by microevolution creating an a stable equilibrium body size distribution. More recently, I have identified a non-trend in the macroevolution of coloniality and photosymbiosis in scleractinian corals (Simpson 2013). In corals, the relative frequencies of both traits have been maintained for 200 million years due to the interaction between species selection for solitary photosymbiotic corals and microevolution favoring solitary/non-photosymbiotic or colonial/photosymbiotic.

Non-trends are phenomena that are impossible to recognize without first understanding how species selection operates. If it turned out that micro and macroevolution only interact additively, non-trends would be rare, only occurring where micro and macroevolutionary processes are equal magnitude and opposite in direction. But micro and macroevolution also interact multiplicatively because microevolutionary changes produce the patterns of species-level variation and heritability; consequently a range of interactions will produce non-trends and may be the most common consequence of species selection in nature.

## Unanswered questions and conclusions

We have learned a considerable amount about the nature of macroevolution since the 1970s. This progress has occurred through an interplay between empirical, conceptual, and theoretical advances. Many conceptual issues about species selection could not be fully resolved until robust methods for quantifying taxonomic rates existed (e.g., Alroy 2000, Foote 2000a, b, 2003). These rate metrics are unbiased by small sample sizes, and because they can be used in a likelihood framework, they can also be used to statistically distinguish between cases with rate differences and those without (Kiessling and Simpson 2010, Simpson and Harnik 2009). With these empirical tools it is possible finally see Arnold and Fristrup’s (1982) ideas grow.

The conceptual framework that Price’s theorem provides is particularly powerful because of its ability to link between theory and empirical application. Its terms and variables are expressed in statistical terms that can be directly measured in the fossil record. I prefer Price’s theorem for thinking about macroevolution for this reason. It speeds up the development of theory, immediately suggests empirical observations to test out new ideas, and helps organize independent observations into coherent results.

Ironically, just as in the study of large-scale trends in the time prior to Simpson’s Tempo and Mode, Price’s theorem is useful because it decomposes the patterns that produce trends into components that consist of a single process. But unlike the state of knowledge in the time before Simpson, Price’s theorem gives us the ability to put the processes back together and study how they interact.

There remain several open empirical issues with species selection. One important one is if organismal and emergent traits differ in their role in species selection. By hitchhiking with intermediary traits, organismal traits can covary with speciation and extinction. Does hitchhiking create less consistent patterns of differential diversification because of the indirect covariance between an organismal trait and species fitness? Few empirical studies of these interactions have been done (see Harnik 2011 for a rare example). In labrid fishes (Alfaro et al. 2009), the emergent sexual dichromatism and organismal presence of the parrotfish pharyngeal jaw both increase relative diversification rates. The two traits also covary with each other. When I measured the partial regressions between diversification rates and the two traits I found differences between their temporal patterns of selectivity (Simpson and Müller 2012). The organismal-level trait show long term changes in magnitude and direction of selection. Emergent sexual dichromatism, however, was always strongly associated with high diversification rates.

This result hints that the intuition some have had about organismal traits and species selection may have some degree of truth to it (Gould and Eldredge 1988, Lieberman and Vrba 1995, Vrba 1983). Even though organismal traits can be seen by species selection, because of hitchhiking, the link can be weak and wobbly. Organismal traits may be prone to variable magnitude and direction of species selection and also to macroevolution by species-level drift. And so emergent traits may show a tendency to have more stable magnitudes and directions of species selection relative to organismal traits.

Because of the weak connection between organismal traits and species selection, it is unlikely that species selection will directly result in adaptations in organisms within species, although it is theoretically possible given the right circumstances (Rice 1995). Rather, the dominant role of species selection in the history of life is to influence the frequencies of traits among species.

Interestingly, species selection may have an important role in the evolution of homologous traits from novelties. Homologous traits share a common “character identity network” but may differ in any other way including form and position (Wagner 2014). Novel identity networks evolve, either by the origin of new recursive gene network components, or by new recursive links between preexisting genes. The ability for novelties to become homologous characters depends on their proliferation in frequency across species (Wagner 2014). An increase in frequency of homologues among species permits their divergence and the subsequent evolution of character states. What this means in macroevolutionary terms is that homologous traits may have to be historically favored by species selection. This potential role of species selection in the evolution and innovation of homologous characters is totally unexplored and an exciting frontier for research integrating macroevolution and the evolution of development.

## Acknowledgements

Thanks to Doug Erwin, Dan McShea, Melanie Hopkins, Chris Haufe, Aaron Bradley, Talia Karim, Sarah Tweedt, and Rachel Warnock for discussion and comments. This work was funded by the Springer fund in the Department of Paleobiology at the Smithsonian Institution’s National Museum of Natural History.

